# Towards label-free non-invasive autofluorescence multispectral imaging for melanoma diagnosis

**DOI:** 10.1101/2023.09.25.559240

**Authors:** Aline Knab, Ayad G. Anwer, Bernadette Pedersen, Shannon Handley, Abhilash Goud Marupally, Abbas Habibalahi, Ewa M. Goldys

**Affiliations:** Graduate School of Biomedical Engineering, Faculty of Engineering, University of New South Wales, Sydney, 2052, Australia; ARC Centre of Excellence for Nanoscale Biophotonics, University of New South Wales, Sydney, 2052, Australia; Macquarie Medical School, Faculty of Medicine, Health and Human Sciences, Macquarie University, Sydney, NSW, Australia; Melanoma Institute Australia, The University of Sydney, Sydney, NSW, Australia

**Author notes:** Corresponding authors: Aline Knab, Ewa Goldys, Abbas Habibalahi, Graduate School of Biomedical Engineering, Faculty of Engineering, University of New South Wales, Sydney, 2052, Australia. Equal senior authors (Abbas Habibalahi, Ewa M. Goldys).

**Keywords:** autofluorescence, multispectral, melanoma, fibroblasts, machine learning, feature analysis, label-free

## Abstract

This study focuses on the use of cellular autofluorescence which visualizes the cell metabolism by monitoring endogenous fluorophores including NAD(P)H and flavins. It explores the potential of multispectral imaging of native fluorophores in melanoma diagnostics using excitation wavelengths ranging from 340 nm to 510 nm and emission wavelengths above 391 nm. Cultured immortalized cells are utilized to compare the autofluorescent signatures of two melanoma cell lines to one fibroblast cell line. Feature analysis identifies the most significant and least correlated features for differentiating the cells. The investigation successfully applies this analysis to pre-processed, noise-removed images and original background-corrupted data. Furthermore, the applicability of distinguishing melanomas and healthy fibroblasts based on their autofluorescent characteristics is validated using patient cells with the same evaluation technique. Additionally, the study tentatively maps the detected features to underlying biological processes. This research demonstrates the potential of cellular autofluorescence as a promising tool for melanoma diagnostics.

## Introduction

Melanomas are the most invasive type of skin cancer and although several effective systemic treatments are now available, mortality rates remain high in patients diagnosed with advanced melanoma [1, 2]. The risk of developing melanoma is influenced by genetic risk factors such as fair skin, light hair color, blue eyes, and freckles coupled with prolonged exposure to sun light and UV radiation [3, 4]. These factors contribute to high incidence rates in Australasia, Europe, and North America which accounted for 275 000 out of the 325 000 reported melanoma cases in 2020 [5, 6]. If diagnosed early, the 5-year survival rate is above 90% [2, 5, 7]. The standard skin screening procedure for skin lesions including melanoma is visual inspection through dermoscopy, and its accuracy depends on the examining physician’s knowledge and experience [8]. In addition, rare melanoma types with atypical features such as hypopigmented amelanotic melanoma are particularly difficult to diagnose with standard methods [9-11]. As a result, some suspicious skin changes may be missed and would not be referred to for follow-up biopsies for a final histological assessment. Therefore, noninvasive accurate imaging technologies for rapid standardized melanoma assessment would be valuable. Optical detection methods such as reflectance confocal microscopy [12], optical coherence tomography [13], multiphoton imaging [14], and stepwise two-photon excited fluorescence [15] have been investigated for melanoma diagnostics. Significant effort has been invested in non-invasive image-based melanoma detection using conventional skin images analyzed by machine learning and artificial intelligence [16, 17]. While these methods improve melanoma detection, issues arise because of costs and varying accuracies [16, 18].

This study explores the feasibility of a non-invasive multispectral imaging approach for melanoma detection. Cell transformation can alter cell metabolism and a common feature is the Warburg effect associated with increased glucose uptake and aerobic glycolysis in cancer cells compared to healthy cells [19, 20]. Cellular autofluorescence reflects metabolic processes because they involve endogenous fluorophores, such as the co-enzymes nicotinamide adenine dinucleotide (NAD(P)H) and flavins [19, 21, 22]. The fluorescence signals of NAD(P)H (excitation maxima at 290 and 351 nm, emission maxima at 440 and 460 nm) and flavins such as flavin adenine dinucleotide (FAD, excitation maximum at 450 nm, emission maximum at 535 nm) has been used to represent metabolic processes and is referred to as the optical redox ratio, here defined as flavins/NAD(P)H [19]. As metabolism tends to be dysregulated in cancer, the autofluorescent signatures and in particular the redox ratio have high potential to be useful in oncology applications [19].

The present study investigates the possible application of autofluorescent signatures in melanoma diagnostics with multispectral imaging to enhance accuracy and provide a detailed spectral analysis. It shows the utility of label-free non-invasive multispectral imaging in distinguishing two melanoma cell lines and fibroblasts. The study design also incorporates rare hypopigmented melanomas in the form of amelanotic melanomas in addition to the prevalent hyperpigmented melanomas [10]. Feature analysis was utilized to read and evaluate the molecular fingerprint provided by the autofluorescent signatures. The clinical applicability of the findings is supported by a demonstration on patient melanoma samples and fibroblasts.

## Method

### 1-Sample preparation

### A-Cell Lines

Fibroblast cell line 142BR (90011806) and melanoma cell lines COLO679 (87061210) and A375 (88113005) were obtained from CellBank Australia (Westmead, NSW, Australia).

142BR cell line was sub cultured and maintained in the complete culture medium Minimum Essential Medium Eagle (EMEM + non-essential amino acids, Sigma Aldrich) combined with 2 mM glutamine (Gibco), 15% fetal bovine serum (FBS, Gibco), and 5 ml penicillin/streptomycin (P/S; 100U/ml; Gibco). The melanoma cells COLO679 were cultured in RPMI 1640 with L-Glutamine (Gibco), 10% FBS, and 5 ml P/S. For A375, DMEM (high glucose, pyruvate, no glutamine; Gibco) was used along with 15% FBS and 5 ml P/S. Cells were incubated at 37°C 5% CO2 incubator. Passaging of cells was performed once the confluency reached 80%. Cells were washed with phosphate buffered saline (PBS) and trypsinized with TrypLE (GIBCO, Australia, Catalog No: 12563-029). Following incubation with trypsin for 5 minutes at 37°C, the complete medium was added to trypsinized cells. The cell suspension was centrifuged at 500 g for 5 minutes. After removing the supernatant, the cell pellet was resuspended in the complete medium. Cell viability testing was performed using Trypan blue 0.4% (Sigma Aldrich, Australia, Catalog No: T8154).

### B-Patient Cells

Clinical samples of 10 patients were received under the permission of the Macquarie University Human Research Ethics Committee, reference number 5201400458. Upon receipt, the patient cells were cultured in Dulbeco’s Modified Eagle Medium containing 10% heat inactivated FBS (Sigma Aldrich), 11.25 mM glutamine (Gibco, Thermo Fisher), and 10 mM HEPES (Gibco) and maintained at 37°C and 5% CO_2_.

### C-Preparing cells for spectral imaging

To perform hyperspectral imaging, the cells were seeded into 35 mm plastic culture dishes with 18 mm well and # 1.5 cover slip bottoms (Cell E&G, USA, and # GDB0004-200). Each dish was seeded with 1 ml of cells (10^5^ cells/ml) and incubated at 37°C, 5% CO_2_ for 48 hours to reach 70% confluence.

The cells were washed three times with PBS prior to imaging, then 4 ml of Hanks Balanced Salt Solution (HBSS) was added into each dish. A total of 6 dishes of 142BR (3xP27, 3xP28, 2xP30), 5 dishes of COLO679 (3xP11, 2xP13, 2xP21), and 5 dishes of A375 (5xP7) were analyzed resulting in a minimum of 300 imaged cells per group.

The same cell preparation protocol was used for imaging patients’ cells.

## 2-Experimental set-up

Multispectral imaging of cellular autofluorescence was carried out using fluorescence microscopes customized to measure cellular autofluorescence designed by Quantitative Pty Ltd. To this aim, regular fluorescence Olympus IX83 microscopes were retrofitted with a multi-LED light source and filter cubes containing excitation, edge, and emission filters as described in [23, 24] and schematically shown in Supplementary Figure 1. A cooled, highly sensitive camera was integrated into the system to capture the autofluorescence signals. The experimental setup allowed the collection of brightfield images and multispectral images captured on a 40x oil objective (NA 1.35).

For immortalized cells, the imaging system was coupled with a Nüvü™ EMCCD camera HNü 1024. The imaging protocol defined a set of 33 spectral channels (see Supplementary Table 1 for a list of channels). The channels covered excitation wavelengths ranging from 345 nm to 505 nm and captured emission wavelengths above 391 nm. To ensure optimal signal-to-noise ratios, exposure times and image averaging were adjusted individually for each channel.

**Table 1:** Specifications of selected feature IDs (FIDs) for immortalized cells with (column 1) and without (column 2) image pre-processing and selected features for patient cells (column 3)

| FID | Immortalized cells,<br>with pre-processing | Immortalized cells,<br>without pre-processing | Patient cells |
| --- | --- | --- | --- |
| 1 | ex 490 nm, em 558-604 nm/<br>ex 371 nm, em 398-504 nm | ex 385 nm, em 558-604 nm/<br>ex 403 nm, em 611-900 nm | ex 391 nm, em 454-495 nm |
| 2 | ex 397 nm, em 665-1200 nm/<br>ex 430 nm, em 665-1200 nm | ex 403 nm, em 558-604 nm/<br>ex 476 nm, em 558-604 nm | ex 368 nm, em 573-613 nm/<br>ex 391 nm, em 454-495 nm |
| 3 | - | ex 345 nm, em 396-437 nm/<br>ex 490 nm, em 398-504 nm | ex 413 nm, em 573-613 nm/<br>ex 373 nm, em 454-495 nm |

For clinical samples, a multispectral microscope with an Andor IXON 885 EMCCD camera was employed. The imaging protocol for clinical samples utilised 38 channels (see Supplementary Table 2). The excitation wavelengths ranged from 340 nm to 510 nm, while the emission filters covered the range of 420 nm to 650 nm.

To account for potential uneven illumination of the field of view, and relate the multispectral images to reference fluorescence values measured by a FluoroMax 4 spectrofluorometer (HORIBA Scientific, Japan), a calibration fluid containing 3.75 μM NADH and 1.24 μM FAD was imaged in addition to cellular samples. Moreover, a water image was used to eliminate background artefacts during image pre-processing, ensuring the accuracy and reliability of the experimental results.

## 3-Image pre-processing

The multispectral images contain a range of artefacts such as cosmic ray-induced spikes, readout noise, Poisson’s noise and background autofluorescence, in addition to the cellular autofluorescent signal. Therefore, these artefacts had to be removed before evaluation. High outlier values were eliminated based on the intensity histogram and replaced by the mean value of the surrounding pixels [25]. Wavelet filtering with hard thresholding removed noise enhancing the image quality [26]. For every channel *i*, equation (1) was computed for each pixel (*x,y*) to receive a flattened and calibrated image *cell*_*cal*_(*x,y, i*)to be smoothed in the subsequent step [26].

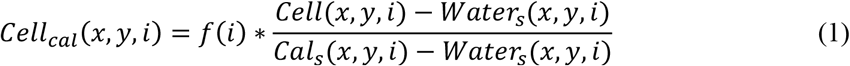

*Water*_*s*_and *cal*_*s*_ denote the respective smoothed water and calibration file and *cell* displays the raw cell file. *f*(*i*)is the calibration factor from the calibrated reference measurement with the sum of all channels normalized to 1. Shifting by the median value of manually defined background areas removed the remaining traces of autofluorescent signatures outside cells.

As this procedure would impede clinical applicability, the experiment on immortalized cells was additionally run without image pre-processing.

## 4-Feature analysis

The feature analysis conducted in this study is directed towards the classification of data into distinct groups of healthy and diseased cells, with a particular focus on identifying different melanoma subtypes when applicable. The analysis was performed on pre-processed images of manually segmented immortalized and patient cells, as well as the original images of immortalized cells. For each cell, the features mean intensity, channel ratios and products were generated. Furthermore, the mean value of the brightest 10% of pixels and statistical measures such as pixel variance, skewness, kurtosis, and entropy were computed, resulting in 924 features for immortalized cell lines and 1634 features for patient cells. To eliminate potential outliers and ensure robust data analysis, the lower and upper one percentile of the overall performance values are removed from the dataset.

The implemented feature selection algorithm with the framework presented in Supplementary Figure 2 effectively reduces exposure times and mitigates overfitting issues. It combines two methods: the ANOVA significance test, which identifies features with *p*< 0.05 as significantly different [27], and the Pearson correlation method which considers r-values between -0.3 and +0.3 as uncorrelated [27, 28]. This study further limits the r-value to the negligible range of -0.1 to +0.1 to ensure feature dissimilarity. The optimal sub-selection of features minimizes the sum of their p-values with the secondary condition of a smaller r-value.

For pre-processed immortalized cells, the two most informative features were selected, and the three most revealing features were chosen for the non-processed immortalized cells and the biologically variable patient cells. A support vector machine (SVM) classifier was trained on the selected features with 60% of the data and validated on the remaining 40% of the dataset using the receiver operating characteristic (ROC) curve and the area under the ROC curve (AUC).

## Results and Discussion

For the immortalized cells 142BR (healthy fibroblasts), COLO679 (melanoma) and A375 (amelanotic melanoma), the algorithm identified two channel ratios as the most informative features (see Table 1). Their feature distribution is displayed in the box plots in Figure 1, c) and d). Figure 1, a) relates the two selected features demonstrating three distinct classes with each data point representing one observation. The ROC curve (Figure 1, b) shows high values for the AUC with 142BR, COLO679, and A375 scoring, respectively 96%, 97%, and 100%.

**Figure 1.**
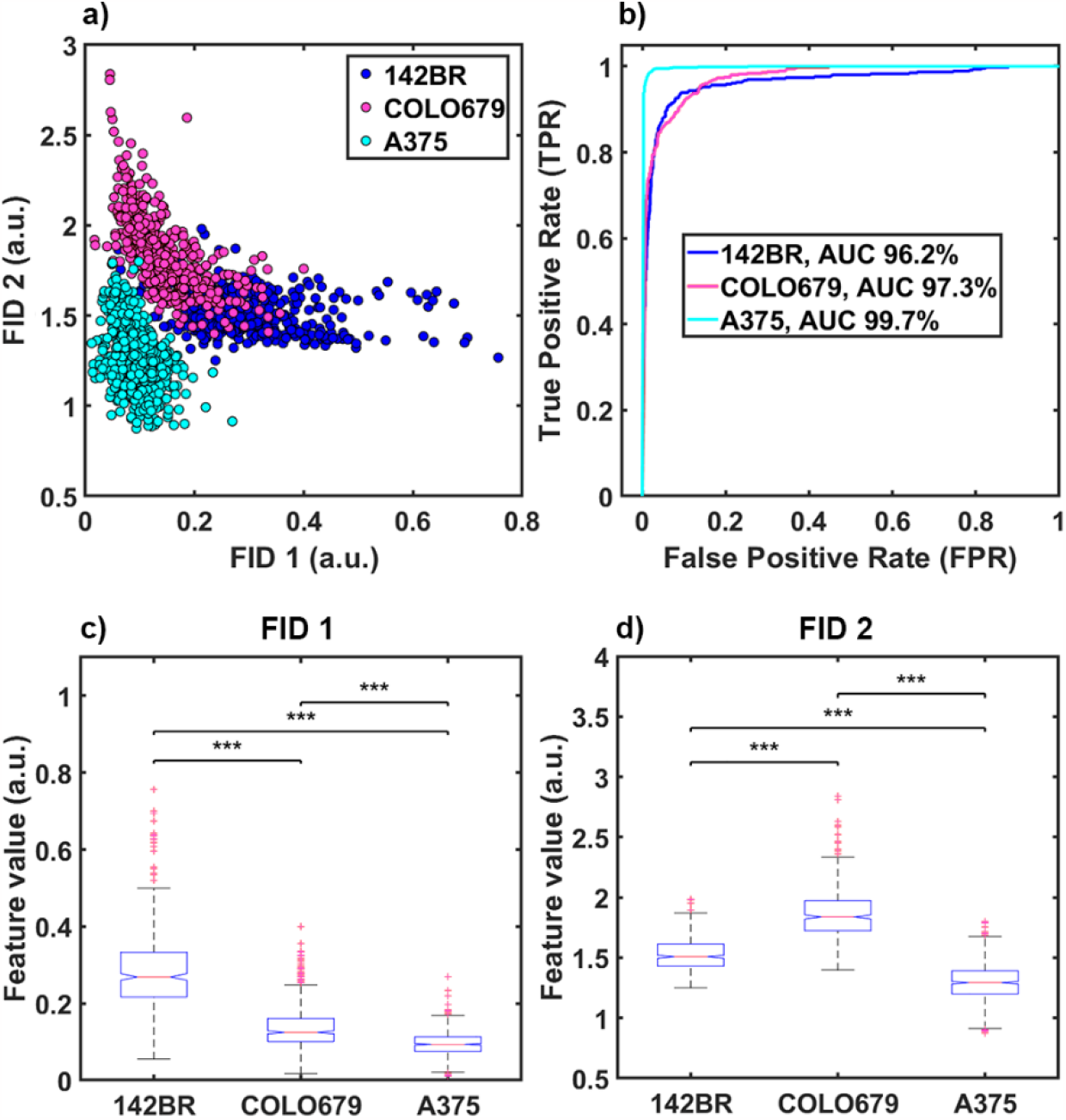
Results of feature analysis after pre-processing for multispectral imaging of healthy fibroblast cells 142BR and the melanoma cell lines COLO679 and A375; (a) 2D feature space; (b) receiver operating characteristics (ROC) and area under the curve (AUC) for classifier evaluation; (c) box plot of feature ID 1; (d) box plot of feature ID 2 (***p<0.0001)

Furthermore, the algorithm was applied to the original multispectral images without image pre-processing. The increased number of outliers in the feature distribution (shown in the boxplots of the three selected features in Figure 2, c) indicates higher data variability than in pre-processed images. The three selected features are plotted in Figure 2, a) forming three separate classes. The predictive power of the classifiers is AUC=89% for 142BR, AUC=85% for COLO679, and AUC=97% for A375.

**Figure 2.**
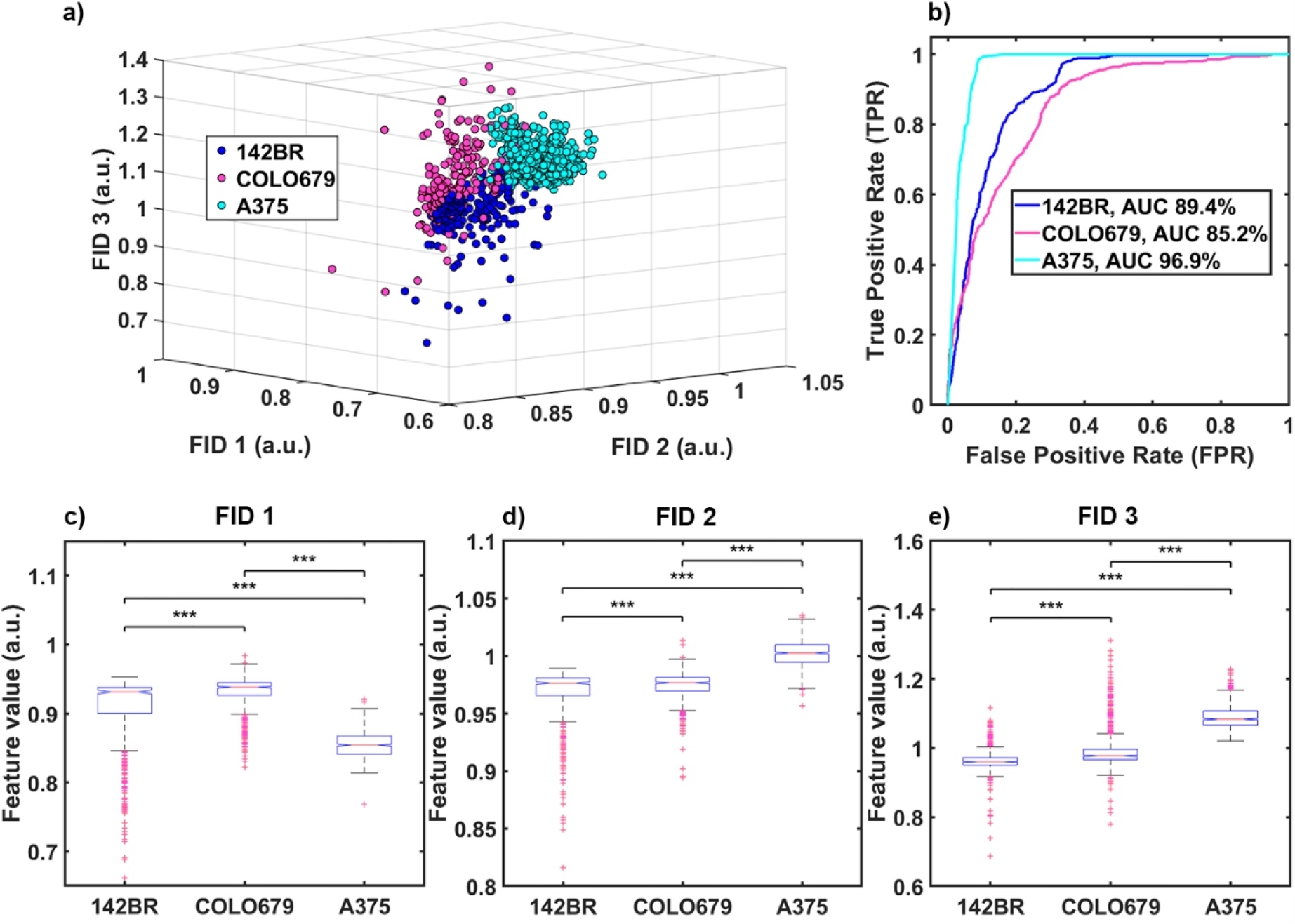
Feature analysis without pre-processing multispectral imaging of healthy fibroblast cells 142BR and the melanoma cell lines COLO679 and A375; (a) 3D feature space; (b) receiver operating characteristics (ROC) and area under the curve (AUC) for classifier evaluation; (c) box plot of feature ID 1; (d) box plot of feature ID 2 (***p<0.0001)

Similar calculations were performed on clinical patient cells, where three distinctive features were selected as excellent representatives, as depicted in the box plots in Figure 3, c)-e). The 3D visualization in Figure 3, a) shows a clear grouping of normal healthy skin cells and melanoma cells. The AUC of 92.8% indicates strong classification accuracy (Figure 3, b).

**Figure 3.**
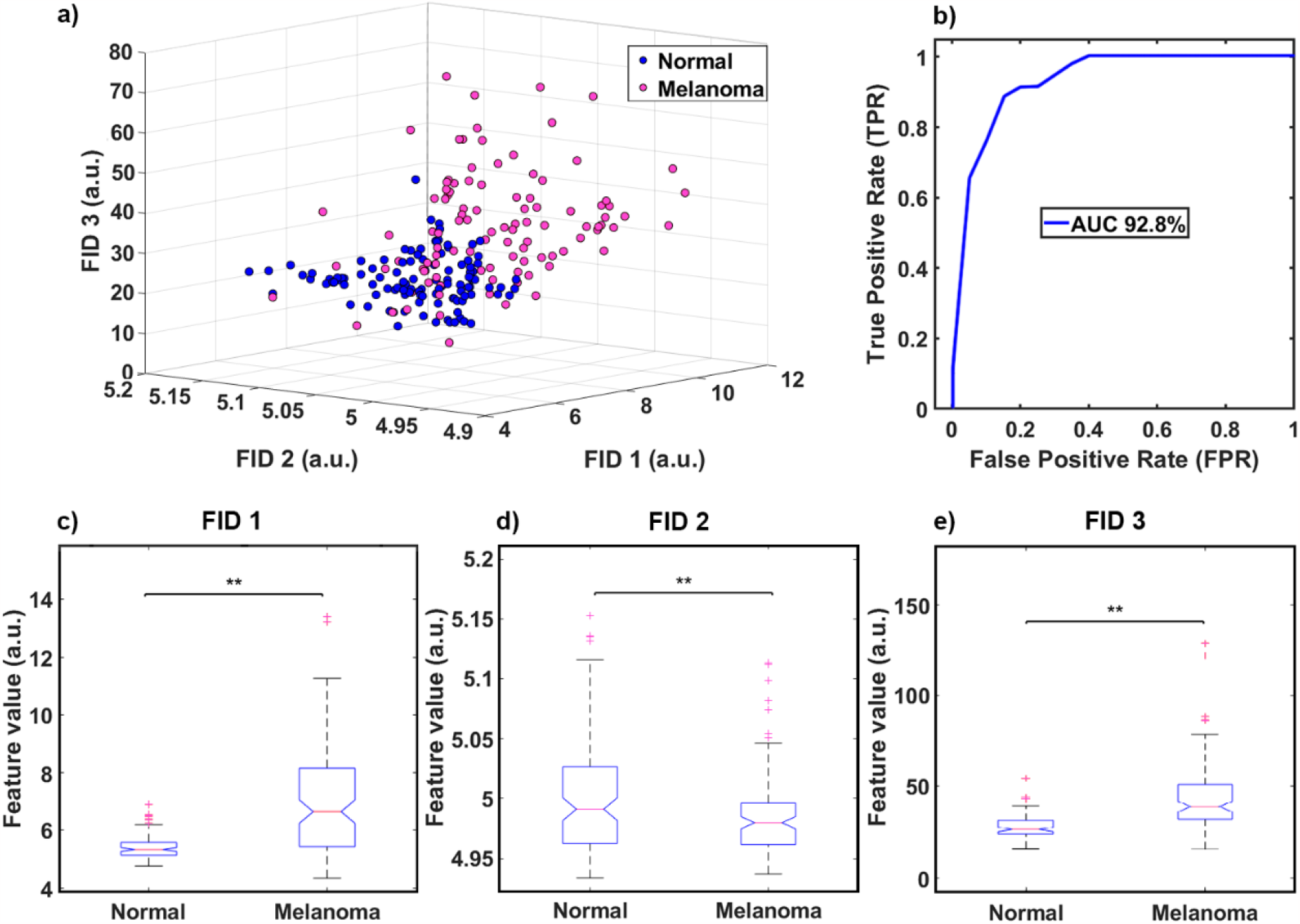
Feature analysis aiming to differentiate clinical melanoma cells from normal fibroblast cells 3D feature space; (b) ROC curve; (c) box plot of feature ID 1; (d) box plot of feature ID 2; (e) box plot of feature ID 3 (**p<0.001)

The measured cellular autofluorescence by multispectral imaging is largely determined by the fluorescence signals NAD(P)H and flavins. The redox ratio (flavins/NAD(P)H) or its inverse is represented as one selected feature in each of the three analysis: For immortalized cells analysed after pre-processing (Figure 1), Feature ID 1 (FID 1) displays flavins/NAD(P)H, whereas the inverse can be found in FID 3 of immortalized cells without pre-processing (Figure 2). For patient cells (Figure 3), the redox ratio is displayed in FID 3 and its inverse in FID 2. Variation in the ratio of these autofluorophores can provide insight into the metabolism of the cells, including altered rates of glucose catabolism, and therefore, the Warburg effect observed in cancer cells [29], having the potential to be used as a marker of early cancer detection. The substantial variation in the optical redox ratio correlates with disparities observed in prior investigations examining melanomas [30].

The experiments on immortalized and patient cells presented here indicate that autofluorescence may be a valuable optical biomarker for melanoma diagnosis. Based on a simple data analysis, distinct groupings of healthy and melanoma cells could be identified using a limited number of features (2 features for pre-processed immortalized cells, 3 features without pre-processing and patient cells). The method can distinguish between healthy skin cells and melanoma cells based on their autofluorescent characteristics on four respective six (immortalized cells) and five spectral channels (patient cells). Limiting channels offers benefits for clinical applicability by reducing the imaging time and data collection, consequently, allowing for a higher throughput and increased patient comfort. Multispectral imaging of cellular autofluorescence achieves a highly accurate differentiation between fibroblasts and two melanoma types on immortalized cell lines both with and without pre-processing. In particular, amelanotic melanomas which are generally detected late due to their hypopigmentation were classified nearly perfectly in pre-processed images. However, accurate image pre-processing steps including background subtraction and noise reduction can hinder clinical applicability. Consequently, we analyzed un-processed images without background subtraction to simulate real-world conditions which also resulted in three clearly separated groups omitting the need of the prior steps. Our evaluation showed that cancer autofluorescence exceeds background autofluorescence from other sources such as the microscope, highlighting its potential as an effective melanoma diagnostic indicator with practical clinical utility. This is reinforced by the technique’s strong data separation in diverse clinical patient samples.

The technology studied in this work has the potential to become a simple handheld diagnostic tool complementing standard dermoscopy. It could easily be integrated with existing imaging methods offering advantageous additional information. Extensive research has focused on image classification for melanoma detection, often with varying accuracies [16, 17]. Combining AI-based melanoma detection with autofluorescent characteristics could enhance early-stage cancer diagnosis and address misclassification challenges. For transferring this technology to human skin tissue, the impact of cell immortalization [31] and potential interference of melanin absorbance across the visible wavelength range [32] when imaging hyperpigmented melanoma must be considered.

Overall, this work shows that label-free non-invasive multispectral imaging of cell autofluorescence could greatly contribute to early-stage melanoma diagnosis by combining regular imaging with insights into cell metabolism.

## Author’s contributions

The study on immortalized cell lines, the study was designed by A.K., A.H., and E.G with assistance of A.A. and A.G. A.K. and A.A. were responsible for cell culturing. A.K. acquired and evaluated the data with the support of S.H and A.G.M. The data was interpreted and discussed by A.K, A.A, A.H, and E.G. The overall study was supervised by A.H. and E.G.

The study of patient cells, it was designed by A.H, H.R, and E.G. A.H. was responsible for acquiring and evaluating the data. The data was interpreted and discussed by A.H, H.R, B.P, A. A, and E.G. Technical, material, and administrative support for the study was provided by H.R, B.P, A. A, A.H, and E.G. The study was supervised by H.R and E.G.

## Acknowledgements

We are thankful for Professor Helen Rizos for her contribution with the clinical aspects and providing clinical samples. Additionally, we also acknowledge contributions from Adnan Agha, Dr. Akanksha Bhargava, Dr. Jared Campbell, and Yuan Tian who provided training, insights, and expertise.

This work was partially supported by the Australian Research Council Centre of Excellence for Nanoscale Biophotonics CE14010003. Aline Knab, Shannon Handley, and Abhilash Goud Marupally acknowledge the PhD scholarship support from University of New South Wales. Dr. Habibalahi is supported by Cancer Institute NSW early career fellowship (2021/ECF1291).

## Conflict of interest

n/a

## Data availability statement

Data which can be accessed without ethical permission is available from the corresponding author upon reasonable request.

## Notes

### Competing Interest Statement

The authors have declared no competing interest.

