## Supplementary Information for "Towards label-free non-invasive autofluorescence multispectral imaging for melanoma diagnosis"

#### Customized microscope for multispectral imaging of cellular autofluorescence

The device set-up is schematically presented in Supplementary Figure 1 for the device used in imaging immortalized cell lines. The customized microscope used in imaging patient samples differs regarding the LED and filter cube selection, and has a different camera attached. While the system for analyzing immortalized cell lines relied on the Nüvü™ EMCCD camera HNü 1024, the device for imaging primary cells works with an Andor IXON 885 EMCCD camera.

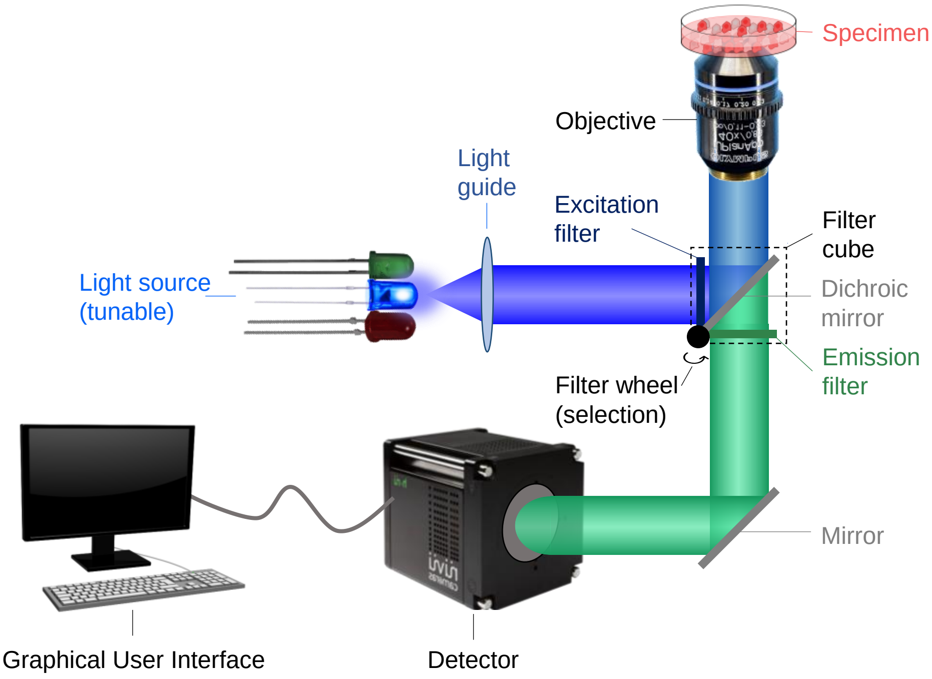


*Supplementary Figure 1: Schematic of the experimental set-up*

### Imaging protocol

Supplementary Table 1 includes the imaging protocol used for analyzing immortalized cell lines (142BR, COLO679, A375). Supplementary Table 2 shows the imaging protocol applied in investigating patient melanoma cells vs. fibroblasts.

| **Immortalized cell lines** | | |
|---|---|---|
| **Channel** | **$\lambda_{ex}$ [nm]** | **$\lambda_{em}$ [nm]** |
| 1 | 345 ± 5 | 396-437 |
| 2 | 358 ± 5 | 396-437 |
| 3 | 371 ± 5 | 396-437 |
| 4 | 345 ± 5 | 398-504 |
| 5 | 358 ± 5 | 398-504 |
| 6 | 371 ± 5 | 398-504 |
| 7 | 345 ± 5 | 558-604 |
| 8 | 490 ± 5 | 558-604 |
| 9 | 505 ± 5 | 558-604 |
| 10 | 385 ± 5 | 558-604 |
| 11 | 397 ± 5 | 558-604 |
| 12 | 403 ± 5 | 558-604 |
| 13 | 418 ± 5 | 558-604 |
| 14 | 437 ± 5 | 558-604 |
| 15 | 451 ± 5 | 558-604 |
| 16 | 476 ± 5 | 558-604 |
| 17 | 345 ± 5 | 611-900 |
| 18 | 490 ± 5 | 611-900 |
| 19 | 371 ± 5 | 611-900 |
| 20 | 385 ± 5 | 611-900 |
| 21 | 403 ± 5 | 611-900 |
| 22 | 418 ± 5 | 611-900 |
| 23 | 437 ± 5 | 611-900 |
| 24 | 469 ± 5 | 611-900 |
| 25 | 397 ± 5 | 665-1200 |
| 26 | 406 ± 5 | 665-1200 |
| 27 | 430 ± 5 | 665-1200 |
| 28 | 457 ± 5 | 665-1200 |

*Supplementary Table 1: Imaging protocol for immortalized cell lines*

| **Patient cells** | | |
|---|---|---|
| **Channel** | **$\lambda_{ex}$ [nm]** | **$\lambda_{em}$ [nm]** |
| 1 | 340 ± 5 | 420-460 |
| 2 | 368 ± 5 | 420-460 |
| 3 | 373 ± 5 | 420-460 |
| 4 | 378 ± 5 | 420-460 |
| 5 | 382 ± 5 | 420-460 |
| 6 | 388 ± 5 | 420-460 |
| 7 | 340 ± 5 | 454-495 |
| 8 | 368 ± 5 | 454-495 |
| 9 | 373 ± 5 | 454-495 |
| 10 | 378 ± 5 | 454-495 |
| 11 | 382 ± 5 | 454-495 |
| 12 | 388 ± 5 | 454-495 |
| 13 | 391 ± 5 | 454-495 |
| 14 | 394 ± 5 | 454-495 |
| 15 | 405 ± 5 | 454-495 |
| 16 | 340 ± 5 | 573-613 |
| 17 | 368 ± 5 | 573-613 |
| 18 | 373 ± 5 | 573-613 |
| 19 | 378 ± 5 | 573-613 |
| 20 | 382 ± 5 | 573-613 |
| 21 | 388 ± 5 | 573-613 |
| 22 | 391 ± 5 | 573-613 |
| 23 | 394 ± 5 | 573-613 |
| 24 | 405 ± 5 | 573-613 |
| 25 | 413 ± 5 | 573-613 |
| 26 | 432 ± 5 | 573-613 |
| 27 | 441 ± 5 | 573-613 |
| 28 | 455 ± 5 | 573-613 |
| 29 | 460 ± 5 | 573-613 |
| 30 | 470 ± 5 | 573-613 |
| 31 | 491 ± 5 | 573-613 |
| 32 | 510 ± 5 | 573-613 |
| 33 | 382 ± 5 | 575-650 |
| 34 | 388 ± 5 | 575-650 |
| 35 | 391 ± 5 | 575-650 |
| 36 | 394 ± 5 | 575-650 |
| 37 | 405 ± 5 | 575-650 |
| 38 | 413 ± 5 | 575-650 |

*Supplementary Table 2: Imaging protocol for patient cells*

### Feature evaluation pipeline

Supplementary Figure 2 illustrates the applied feature evaluation framework. First a feature bank is generated, which includes features such as mean intensity, channel ratios and products, mean value of the brightest 10% of pixels, pixel variance, skewness, kurtosis, and entropy. These features are computed on a single-cell basis. Feature significance is estimated using ANOVA, and features with a p-value above 0.05 are removed. Subsequently, indicative features are selected, with the primary criterion being the minimization of the sum of p-values among the feature sub-selection and a smaller r-value as a secondary condition. The data is divided into 60% training data for constructing a model using an SVM classifier, and the model's performance is validated using the remaining 40% of the dataset.

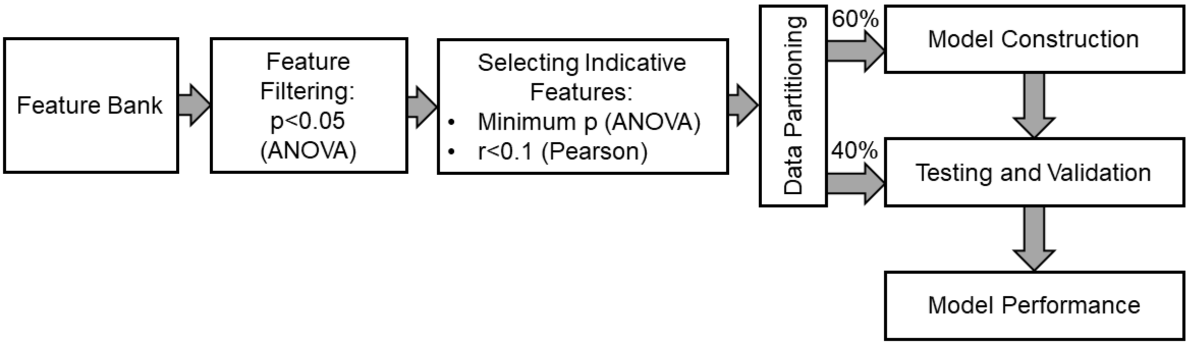


*Supplementary Figure 2: Feature evaluation framework*
